# Utility of Cellular Measurements of Non-Specific Endocytosis to Assess the Target-Independent Clearance of Monoclonal Antibodies

**DOI:** 10.1101/2024.04.16.589824

**Authors:** Mark A. Bryniarski, Md Tariqul Haque Tuhin, Carolyn D. Shomin, Fatemeh Nasrollahi, Eunkyung Clare Ko, Marcus Soto, Kyu Chung, Carrie Poon-Andersen, Ronya Primack, Diana Wong, Esperanza Ojeda, John Chung, Kevin D. Cook, Kip P. Conner

## Abstract

Past studies have demonstrated higher clearance for monoclonal antibodies possessing increased rates of non-specific endocytosis. However, this metric is oftentimes evaluated indirectly using biophysical techniques or cell surface binding studies that may not provide insight into the specific rates of cellular turnover. Furthermore, few examples evaluating non-specific endocytosis have been reported for a therapeutic antibody that reached clinical assessment. In the current report, we evaluated a therapeutic human immunoglobulin G2 monoclonal antibody targeted against the interleukin-4 receptor alpha chain (IL-4Rα) that exhibited elevated target independent clearance in previous Phase 1 and 2 studies. We confirmed high non-specific clearance of the anti-IL-4Rα antibody as compared to a reference antibody during pharmacokinetic assessments in wild type mice where target-mediated disposition was absent. We then developed a cell-based method capable of measuring cellular protein endocytosis and demonstrated the anti-IL-4Rα antibody exhibited marked non-specific uptake relative to the reference compound. Antibody homology modeling identified the anti-IL-4Rα antibody possessed positive charge patches whose removal via targeted mutations substantially reduced its non-specific endocytosis. We then expanded the scope of the study by evaluating a panel of consisting of both preclinical and clinical monoclonal antibodies and demonstrate those with the highest rates of non-specific uptake *in vitro* exhibit elevated target independent clearance, low subcutaneous bioavailability, or both. Our results support the observation that high non-specific endocytosis is a negative attribute in monoclonal antibody development and demonstrate the utility of a generic cell-based screen as a quantitative tool to measure non-specific endocytosis of protein therapeutics at the single-cell level.

**Highlights:**

- Developed a novel, reproducible cellular assay to directly quantify non-specific endocytosis of therapeutic proteins.
- A previous clinical candidate monoclonal antibody with rapid target-independent clearance in mice and humans possessed extensive non-specific endocytosis that was due to exposed positive charge features.
- Demonstration of distinct rates of endocytosis into mammalian cells for disparate monoclonal antibodies, even those with common specificity for targets or isoelectric points.
- Cell-based assay to quantify the potential impact of non-specific endocytosis on target-independent clearance and/or subcutaneous bioavailability of monoclonal antibodies.

## Introduction

Monoclonal antibody (mAb) disposition is generally dictated by both target-dependent and target-independent mechanisms. Multiple factors influence the relative contribution of non-target mediated clearance to overall mAb clearance, which can vary widely across mAbs. These include the development of anti-drug antibodies (1, 2), or a higher propensity for nonspecific endocytosis (NSE) that results in fast target-independent mAb clearance, CL_ind_ (also referred to as non-specific clearance, or linear clearance based on its mathematical descriptions). Accelerated CL_ind_ of a mAb can result from a high propensity for NSE into non-target cell populations as determined by the unique physicochemical properties of each mAb (3–5). In the absence of receptor-mediated endocytosis, mAb NSE can be facilitated by two distinct, concurrent processes: fluid-phase uptake and non-specific adsorptive endocytosis (1, 2, 6–8). Fluid-phase endocytosis (oftentimes referred to as pinocytosis) is an essential cellular process whereby, under basal conditions, extracellular fluid and its contents are internalized by the cell in a constitutively active, yet non-specific manner (8–10). Non-specific adsorptive endocytosis occurs when a protein non-specifically interacts with the cell membrane, such as charge-based attraction, that results in its internalization. Unlike fluid-phase uptake, non-specific adsorptive endocytosis is protein-dependent and can be influenced by factors that includes localized charge (8, 9, 11, 12).

High rates of mAb NSE can lead to an increased likelihood of intracellular catabolism (5). This is true despite pH-dependent mAb recycling by the neonatal Fc receptor (FcRn) because this salvage pathway is not completely efficient. Therefore, a mAb exhibiting more frequent cellular internalization can be degraded at a significantly higher rate within lysosomes than a mAb with relatively lower intracellular exposure (7). Thus, it is critically important during preclinical selection of therapeutic mAb candidates to identify those with high non-specific cellular uptake due to an increased propensity to exhibit poor pharmacokinetics (PK).

Several biophysical techniques have been developed to study the potential for mAbs to undergo NSE. These include baculovirus particle binding, and heparin chromatography (3, 13). However, non-specific mAb internalization measured directly in mammalian cells is rarely assessed. This type of cellular data could be useful for identifying overt PK liabilities early in preclinical drug development in addition to its potential to inform mechanistic PK models for human translation. For instance, computational models that incorporate cellular internalization rates as parameters to predict mAb partitioning from the blood into tissues could benefit from experimental data obtained from a physiologically relevant cell-based platform to provide greater insight into CL_ind_ mechanisms (6). Therefore, the primary aim of the current work was to develop a reproducible and robust cell-based method to quantify mAb NSE.

Our initial work focused on two antibodies: a non-targeting anti-streptavidin fully human immunoglobulin G1 (hIgG1) mAb (ASA_hIgG1-WT_) as a control for ‘typical’ behavior, and a hIgG2 mAb targeted against the alpha chain of human interleukin-4 receptor (anti-IL-4Rα mAb). The anti-IL-4Rα mAb was examined retrospectively because it exhibited a high CL_ind_ of 11 mL/d/kg and poor subcutaneous bioavailability (F_SC_, 24.3%) in clinical trials, and thus represented an ideal therapeutic candidate for which to test our platform (14). Its clearance can be categorized as high when compared to a panel of 64 cross-industry antibodies in which CL_ind_ was estimated using a statistical modelling approach (15).

First, we confirmed a higher rate of CL_ind_ for the anti-IL-4Rα mAb relative to ASA_hIgG1-WT_ via single dose PK studies in human FcRn transgenic mice. This was followed by the development and application of a novel cell-based method that demonstrated substantially increased NSE of the anti-IL-4Rα mAb when compared to ASA_hIgG1-WT_. Next, we established an experimental workflow to accurately quantify cellular endocytosis across experimental days and confirmed that exposed surface charge on the anti-IL-4Rα mAb was driving its high rate of NSE and rapid CL_ind_. Finally, we applied our established assay workflow to characterize NSE across a diverse set of preclinical and clinical mAbs to validate our assumptions of a direct relationship between NSE-driven cellular endocytosis and *in vivo* PK parameters. Our results demonstrate the utility of a convenient cell-based assessment of mAb NSE for evaluation of potential off-target mAb PK liabilities, and more broadly demonstrate the potential of standardized *in vitro* ADME workflows to support preclinical development of therapeutic mAbs.

## Experimental Section

### Cell Culture

Parental Chinese Hamster Ovary-K1 (CHO-K1) cells were obtained internally from Amgen and maintained in Ham’s F-12K media (Thermo Fisher, 21127022) containing 10% heat inactivated fetal bovine serum (Thermo, 16140071). Vero cells were purchased from the American Type Culture Collection (ATCC; CCL-81, lot 70016956) and grown in ATCC-formulated EMEM with 10% heat-inactivated fetal bovine serum (ATCC; 30-2003). Cells were cultured without antibiotics in a humidified, 5% CO_2_ incubator at 37°C and passaged utilizing 0.05% trypsin-EDTA (Thermo, 25300054). Mycoplasma testing was conducted using the MycoAlert Detection Kit (Lonza). Cells were cryopreserved using complete growth media containing 5% (*v/v*) DMSO.

### Generation of mAbs

Two fully human, wild type mAbs were used for our initial characterization: an anti-IL-4Rα mAb (human IgG2; hIgG2) and a human IgG1 (hIgG1) control mAb raised against streptavidin (anti-streptavidin antibody, ASA_hIgG1-WT_). Both mAbs along with the anti-IL-4Rα mAb mutant series and an ASA hIgG2 mAb were constructed by recombinant DNA technology and produced in stably transfected CHO cells using standardized protocols. The preclinical mAb panel included five antibodies raised against a human target that were anticipated to have negligible cross-reactivity with murine antigen (mAbs B1-5). The clinical panel comprised (1) the drug products of rituximab, pembrolizumab, and daratumumab, (2) research analogs of ustekinumab, bococizumab, infliximab, bimagrumab, sirukumab, duligotuzumab, briakinumab, basiliximab, and bavituximab (referred to as mAb-SEFL2), and (3) 5 internal Amgen mAbs (including the wild type anti-IL-4Rα mAb). The anti-IL-4Rα WT and mutants, two of the three ASA mAbs, and the internal Amgen mAbs were wild type IgGs (i.e. wild type human Fc regions). The ASA_hIgG1-SEFL2,_ the preclinical panel B, and the research analogs were produced in stably transfected CHO cells on the Amgen patented stable effector functionless Fc backbone (SEFL2) as described previously (16).

### PK Studies and Analyses

PK data was obtained from three separate studies previously conducted at Amgen, Inc. to reduce animal usage: (1) ASA_hIgG1-WT_, (2) wild type (WT) anti-IL-4Rα mAb and the EEES anti-IL-4Rα mAb mutant (EEES refers to the abbreviation of the conducted amino acid substitutions; fully described below), and (3) the preclinical mAb panel B. All rodents were purchased from Jackson Laboratory (Bar Harbor, MA). Male and female homozygous Tg276 mice were used to characterize the disposition of ASA_hIgG1-WT_, WT anti-IL-4Rα mAb, and the EEES anti-IL-4Rα mAb mutant. Female BALB/c mice were used for the preclinical mAb panel B. The proteins of interest were administered as a 1 mg/kg (ASA_hIgG1-WT_, WT and EEES anti-IL-4Rα mAbs studies) or 2 mg/kg (preclinical panel B study) intravenous bolus dose via the lateral tail vein. Blood specimens were collected at various times post injection, incubated at ambient temperature for approximately 20 minutes or until fully clotted, and then centrifuged to separate the serum. All serum specimens were stored at −70 °C (± 10 °C) until use in analytical assays. Mice were cared for in accordance with the Guide for the Care and Use of Laboratory Animals, 8th Edition at AAALAC, International accredited facilities. All mice protocols were approved by the Amgen, Inc. Institutional Animal Care and Use Committee (Thousand Oaks, CA). Quantitation of proteins in mouse serum was performed via electro-chemiluminescent immunoassays on the MSD Sector 600 instrument (Meso Scale Diagnostics, MSD; Rockville, MD) using the Amgen clone 35 anti-human Fc antibody (Ab35) as both the capture and detection reagent. In all assays, the analyte serum concentrations were interpolated from standard curves using the corresponding analyte prepared in pooled mouse serum using Watson LIMS software (Thermo). The murine PK of all tested mAbs was described using a noncompartmental analysis in Phoenix v8 (Certara, Princeton, NJ).

### Detection of Anti-drug Antibodies

The single-dose study examining the disposition of the WT and EEES anti-IL-4Rα mAbs included additional groups and test articles that exhibited dramatic decreases in the terminal slopes of some of the concentration-time profiles (data not shown). This observation led us to postulate that immunogenicity may have been present. For this reason, only the WT and EEES samples were subjected to a homogeneous bridging MSD-based assay to measure the presence of anti-drug antibodies (ADAs). ADAs were captured in solution by combination of biotin-labeled and ruthenium-labeled forms of a test article. Formed complexes were subsequently detected and quantified by electrochemiluminescence in a streptavidin-coated MSD plate. Therefore, the matching capture and detection reagents must be used to measure ADAs against each test article.

Mouse anti-hIgG mAb (clone 35) was used as a non-drug specific positive control antibody that was prepared at high and low concentrations, e.g. 10 and 0.2 μg/mL respectively. Multiple naïve individual mouse serum samples were included in each assay plate to determine a plate-specific screening cutpoint (SCP). The SCP defines the threshold between screening-positive and -negative signals and was determined by adding 5x standard deviation to a mean electrochemiluminescence value generated from the naïve serum samples. Any study samples that resulted in an electrochemiluminescence value higher than the SCP were considered ADA-positive. Naïve pooled mouse serum sample was also included in the assay as a negative control.

Samples were pre-diluted 10-fold in assay buffer (low cross buffer #100500, CANDOR Bioscience, Germany). The diluted samples were then mixed with the same volume of reaction mixture composed of 1 μg/mL of biotin-labeled test article and 1 μg/mL of ruthenium-labeled test article in the assay buffer. The mixture was incubated for 2 hours at room temperature with shaking at 600 rpm. Meanwhile, the MSD assay plate (Gold 96-well Streptavidin plate, Meso Scale Diagnostics #L15SA) was added with 150 μL/well of blocking buffer (SuperBlock T20 (PBS) Blocking buffer, Thermo # 37516) and incubated for 2 hours at room temperature without shaking. The blocked plate was then washed three times with 350 μL/well of washing buffer. The sample-reaction mixtures from the incubation were transferred to the blocked assay plate (50 μL/well) and incubated for 1 hour at RT with shaking at 600 rpm. The plate was washed three times with 350 μL/well of the washing buffer. Reading buffer (1X, diluted with ddH_2_O from the stock; 150 μL/well) was added (MSD Read Buffer T (4X) with surfactant, Meso Scale Diagnostics #R92TC) and the plate was read using an MSD reader.

### NSE Studies with Immunostaining and Flow Cytometry

CHO-K1 and Vero cells were seeded onto 96-well plates at a density of 150,000 cells per well 24 h prior to experimentation, or 75,000 cells per well 48 h beforehand. On study days, cells were washed twice in either pre-warmed (37°C) or ice-cold (4°C) Ringer’s solution (pH 7.4, 122.5 mM NaCl, 5.4 mM KCl, 1.2 mM CaCl2, 0.8 mM MgCl2, 0.8 mM Na2HPO4, 0.2 mM NaH2PO4, 5.5 mM d-glucose, and 10 mM HEPES). Fresh Ringer’s solution was added a third time and cells were equilibrated at either temperature for 30 min. Afterwards, wells were aspirated, and then the treatments were immediately applied to cells. Studies utilized 37°C (enables cell surface binding, internalization, intracellular trafficking) and 4°C (enables cell surface binding) conditions. For the time-dependent study, CHO-K1 cells were incubated with 100 µg/mL mAb for 15, 30, 60, or 120 mins. For the concentration-dependent study, CHO-K1 cells received concentrations diluted 1:2 spanning 25 to 200 µg/mL for 60 min. These conditions were based on preliminary data that were found to lie below the limit of detection saturation. For the other mAb assessments, CHO-K1 and Vero cells were incubated with mAbs for 60 min at 100 µg/mL. Following incubations, cells were washed four times on ice with ice-cold Ringer’s, then trypsinized for 4-6 min at 37°C. Once cells were detached, complete ice-cold growth media was added at a 1:1 ratio (*v/v*) to inhibit the trypsin. Cells were removed from cell culture plates and placed into V-bottom 96-well plates (Fisher #249944), then centrifuged at 300 g for 5 min at 4°C. Cells were stained in 100 µL 1X PBS containing 0.5% (*v/v*) Zombie UV fixable viability dye (BioLegend # 423108) on ice for 20 min. Cells were washed once with 100 µL FACS buffer (1X PBS, 2% w/v BSA, 1mM EDTA, 0.1% w/v sodium azide), centrifuged at 300 g for 5 min at 4°C, then fixed/permeabilized in the dark for 20 min at room temperature using the Cyto-Fast fix/perm buffer (Biolegend # 426803). Cells were then washed twice using 1X Cyto-Fast Perm Wash solution (Biolegend # 426803). Samples were next incubated with 5 µg/mL (100 µL per well, Cyto-Fast Perm Wash solution) of an Amgen mouse anti-human Fc monoclonal IgG1 antibody fluorescently conjugated with Alexa Fluor 647 (Ab35-AF647) for 30 min on ice in the dark (17). Samples were washed thrice with Cyto-Fast Perm Wash solution, resuspended in 100 µL Cyto-Fast Perm Wash solution, and analyzed on a BD FACSymphony flow cytometer with an 18 color, 5-laser configuration (UV-355 nm, violet-405 nm, blue-488 nm, yellow/green-561 nm, red-637 nm) and a BD Biosciences High Throughput Sampler, (Catalog #338301). Gating was conducted as depicted within figures, obtaining a target of 10,000 single-cell, Zombie^-^(live) events using BD Diva software. For Vero cell samples, approximately 5000 single live cell events were captured due to differences in trypsin dissociation rates between Vero and CHO-K1 cell lines, which led to less Vero cells being collected. Data was analyzed using FlowJo software (Becton Dickson, Franklin Lakes, NJ) to obtain median fluorescent intensities of the indicated cell populations.

### Antibody Binding Capacity (ABC)

Quantum™ Simply Cellular® Mouse IgG beads were obtained from Bio-Rad (#FCSC815; lot #15515; Hercules, CA). The initial evaluation utilized final Ab35-AF647 concentrations of 5 or 10 µg/mL exactly following the vendor product information with all procedures done on ice or at 4°C. Subsequent work used a concentration of 5 µg/mL to match cell staining experiments as we demonstrated this concentration saturated each bead population with antibody (see Results). For each uptake study, freshly stained beads were analyzed on the same flow cytometer using identical instrument parameters for the cell samples from that day. Bead median fluorescent intensities were obtained using FlowJo software, then plotted against the known antibody binding capacities per bead population supplied by the vendor via GraphPad Prism. The data was fit to a straight line using simple linear regression. The ABC for each cell sample was then interpolated from the cell median fluorescent intensities. For the clinical mAb panel results, mean ABC values for the untreated samples from that experiment day were subtracted from each sample.

### Antibody NSE with Fluorescence Microscopy

Endocytosis studies in CHO-K1 cells were carried out as described above. Following the washes with ice-cold Ringer’s solution, cells were fixed in Cyto-Fast fix/perm buffer for 20 min at room temperature, and then washed twice with 1X PBS. Samples were then blocked and permeabilized in 1X Cyto-Fast Perm Wash buffer for one hour at room temperature. After aspiration, wells were stained with 1.0 µg/mL Ab35-AF647, 1 µg/mL Hoechst (Thermo #H1399), and 2 µg/mL HCS Cell Mask Blue (Thermo # H32720) in 1X Cyto-Fast Perm Wash buffer for 1 hr at room temperature. Next, samples were washed three times with room temperature 1X PBS and imaged on an Opera Phenix High Content Screening system (PerkinElmer) using a 40x water objective with Z-stack acquisition. Maximum projections were obtained using Columbus software (PerkinElmer). Mean fluorescence intensity measurements for the anti-human Fc staining (Ab35-AF647) were obtained with Columbus by segmenting cells via thresholds for nuclei and cytoplasmic regions to demarcate individual cells. Data depicted in Figures 2 and 6 were obtained using separate microscope settings, which resulted in different extents of fluorescent intensities between untreated and the anti-IL-4Rα mAb.

**Figure 1:**
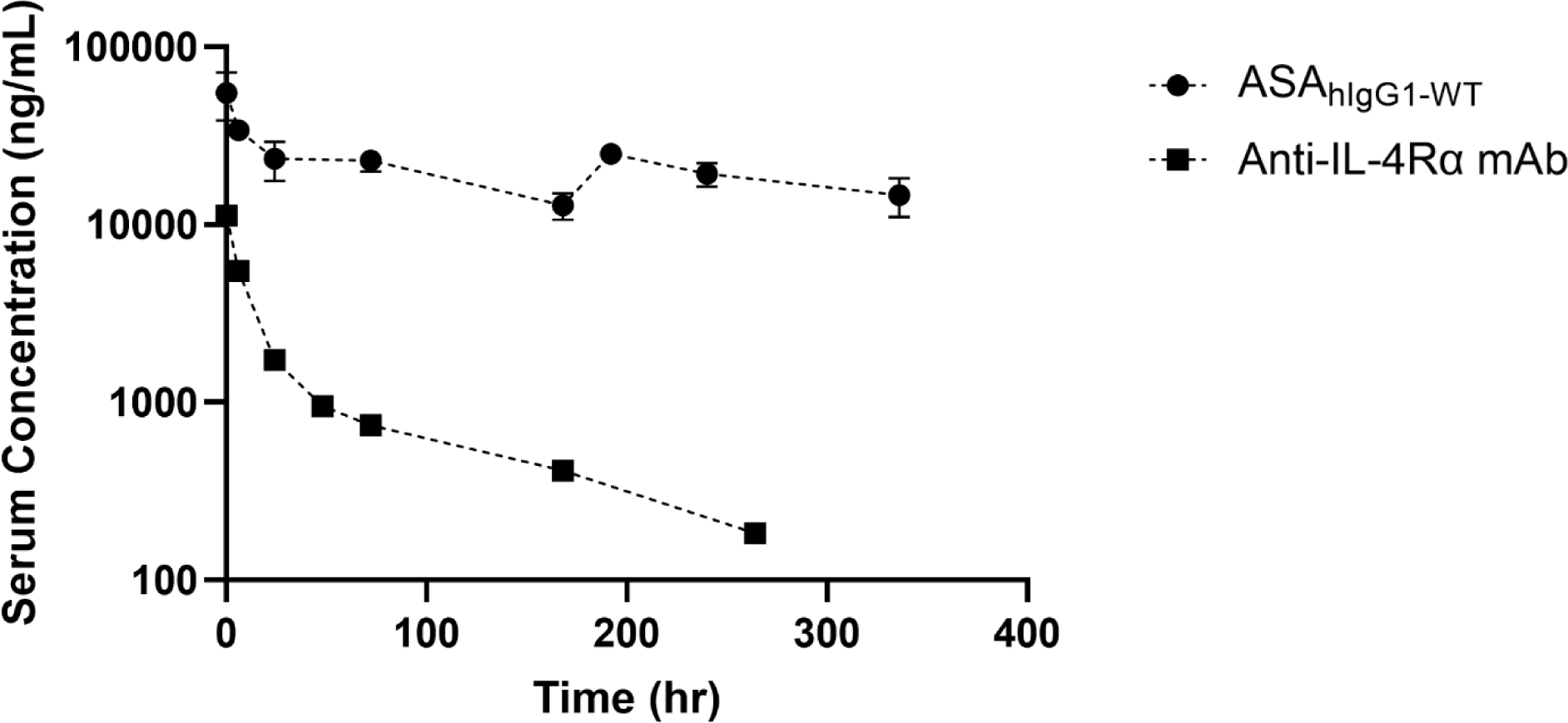
Serum concentration-time profiles of ASA_hIgG1-WT_ and an anti-IL-4Rα mAb in Tg276 mice following 1 mg/kg intravenous bolus doses. The ASA_hIgG1-WT_ possessed substantially lower CL when compared to the anti-IL4Rα mAb. N = 3 separate animals for ASA_hIgG1-WT_, and N = 2 for the anti-IL4Rα mAb due to the detection of anti-drug antibodies in one animal at 264 h.

**Figure 2:**
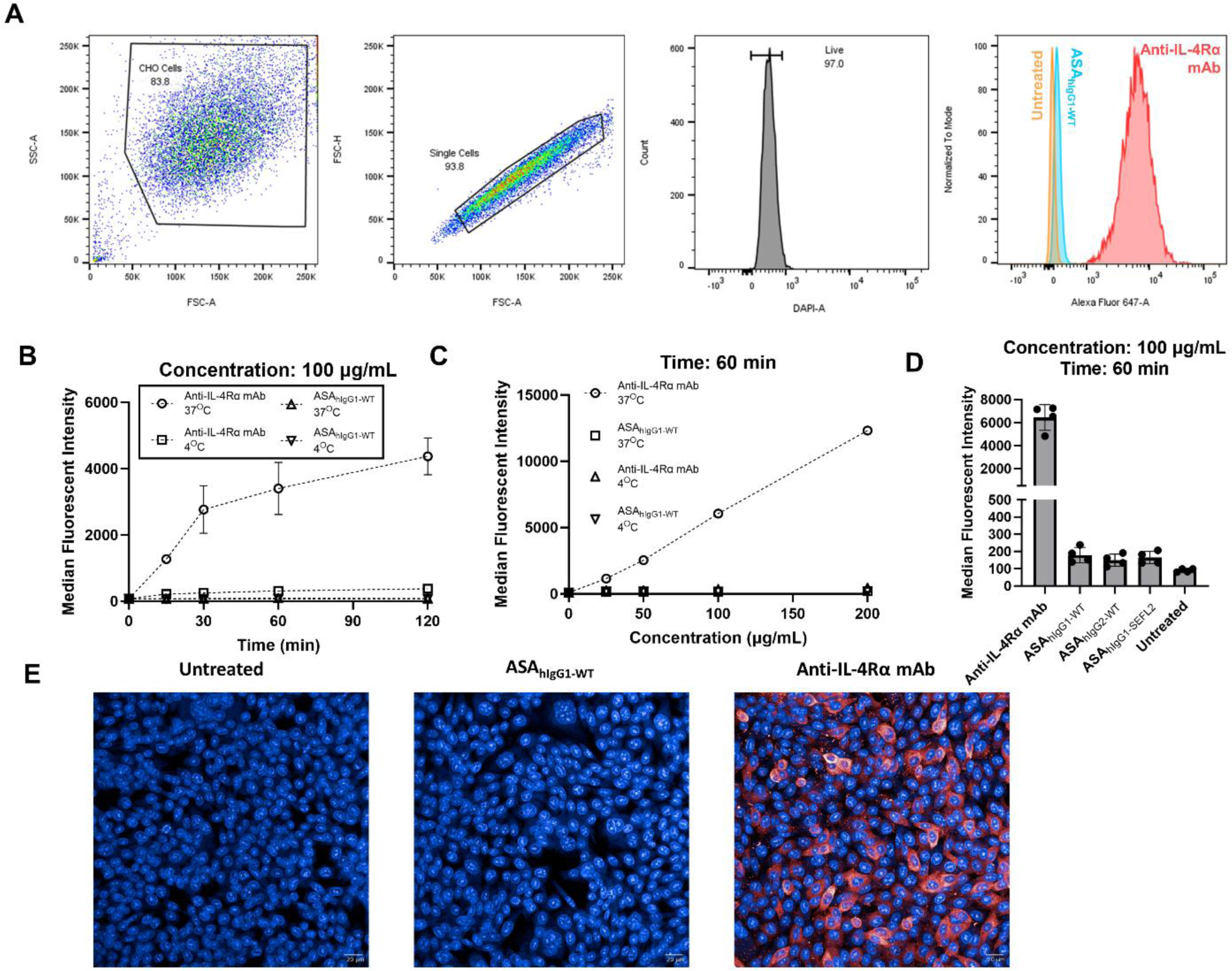
Internalization of the anti-IL-4Rα mAb and ASA_hIgG1-WT_ in CHO-K1 cells. **A** Representative gating strategy used throughout the current study. The median fluorescent intensities of single live cell events were obtained for each sample. **B, C** CHO-K1 cells were incubated with 100 µg/mL anti-IL-4Rα mAb or ASA_hIgG1-WT_ for increasing periods of time at 37°C or 4°C. Biphasic time-dependent (**B**) and linear concentration-dependent (**C**) internalization was observed for the anti-IL-4Rα mAb, but only under 37°C conditions. In contrast, ASA_hIgG1-WT_ endocytosis was negligible at 37°C. Minimal cell surface binding was detected for the anti-IL-4Rα mAb at 4°C relative to the difference between 37°C samples and untreated controls. Monoclonal antibodies were detected in fixed, permeabilized CHO-K1 cells using an Alexa Fluor 647-conjugated anti-human Fc mouse antibody. **D** Uptake studies using three distinct variants of ASA (hIgG1, wild type Fc; hIgG2, wild type Fc; hIgG1, stable effectorless Fc, SEFL2) demonstrated no statistically significant difference (one-way ANOVA with Tukey’s multiple comparisons test) in the amount of non-specific endocytosis under the tested conditions. **E** Immunofluorescence with confocal microscopy was performed following an uptake study at 37°C for 60 min at 100 µg/mL per mAb to confirm the flow cytometry results. Detection was accomplished under fixed, permeabilized conditions utilizing the same anti-human Fc secondary antibody as in **B** and **C**. ASA_hIgG1-WT_, wild type Fc human IgG1 anti-streptavidin antibody; CHO-K1, Chinese Hamster Ovary cells; FSC, forward scatter; SSC, side scatter; -A, parameter area; -H, parameter height.

**Figure 3:**
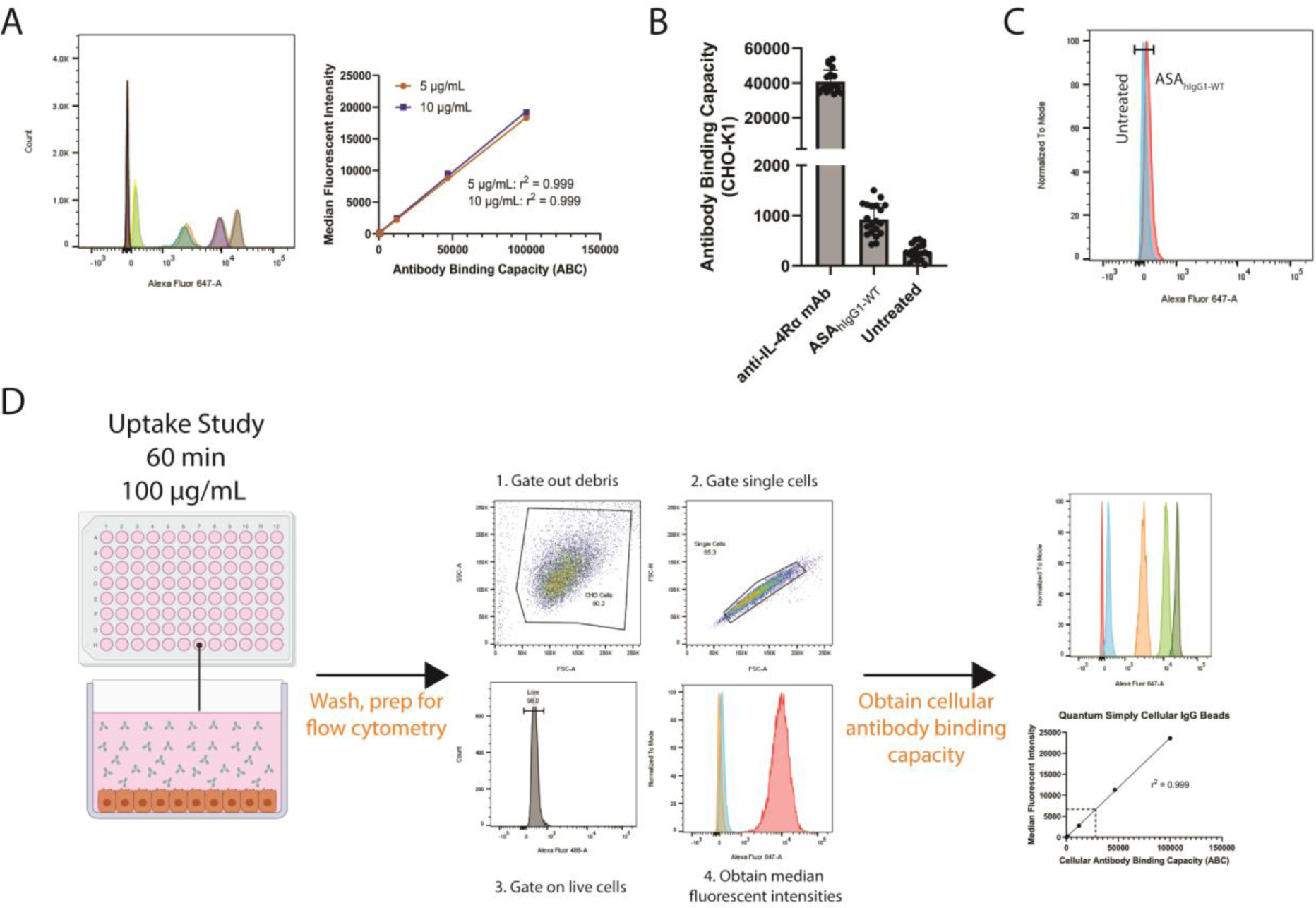
Development of a quantitative endocytosis assay using flow cytometry. **A** Histograms of the five bead populations overlaid when stained with either 5 or 10 µg/mL of the anti-human Fc mouse IgG1 conjugated with Alexa Fluor 647. Plots of the median fluorescent intensities versus the antibody binding capacities (ABCs) indicated both staining conditions resulted in linear curves with r^2^ values of 0.999. **B** Assay reproducibility was evaluated by conducting uptake studies in CHO-K1 cells using 100 µg/mL of anti-IL-4Rα mAb or ASA_hIgG1-WT_ incubated at 37°C for 60 min over several different days. Mean anti-IL-4Rα mAb ABC: 40,839 (SD 6614, CV 16.2%); mean ASA_hIgG1-WT_ ABC: 921 (SD 315, CV 34.2%). N = 20 per sample group. **C** Representative histogram of single ASA_hIgG1-WT_ and untreated samples with a gate incorporating 99% of the untreated population, corresponding to the ‘negative’ signal. The ASA_hIgG1-WT_ endocytosis at the experimental settings was so low that ∼92% of the ASA_hIgG1-WT_ signal resided within the ‘negative’ gate. **D** Illustrative overview of the quantitative endocytosis study. A portion of Figure 3D was generated using BioRender.com. ASA_hIgG1-WT_, wild type Fc human IgG1 anti-streptavidin antibody; IgG1, immunoglobulin G1; CHO-K1, Chinese Hamster Ovary cells; SD, standard deviation; CV, coefficient of variation.

**Figure 4:**
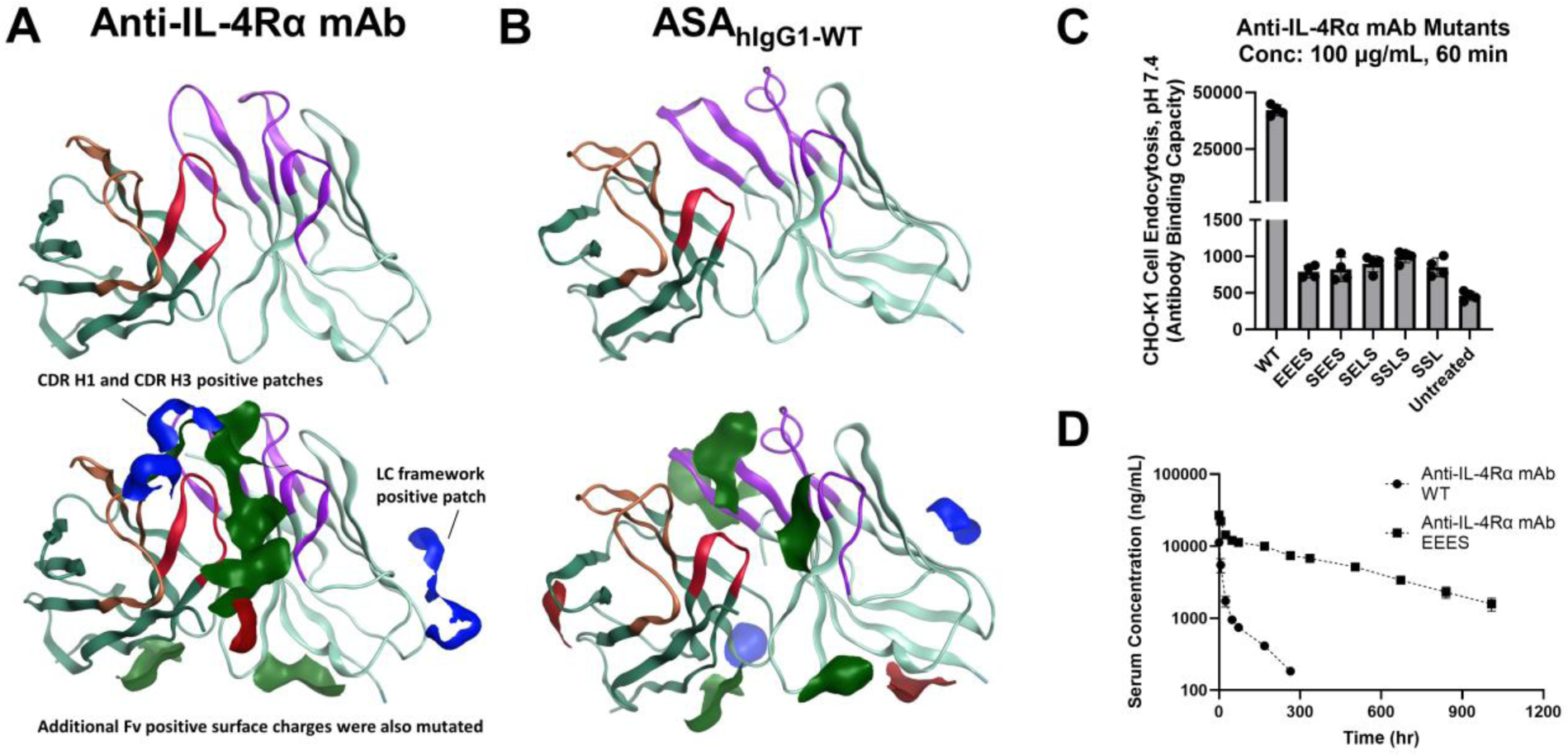
Antibody homology models of the anti-IL-4Rα mAb and ASA_hIgG1-WT_. Fv model ribbon representation (heavy chain in dark teal, light chain in pale teal) depicts heavy chain CDRs in orange (CDR1, CDR2) and red (CDR3). Light chain CDRs are shown in purple. Hydrophobic patches are shown in green. Negative and positive patches are red and blue, respectively. The anti-IL-4Rα mAb (**A**) possessed more positive patches than ASA_hIgG1-WT_ (**B**), with 2 positive patches in its heavy chain CDRs. (**C**) Various point mutations were performed on the anti-IL-4Rα mAb CDR and Fv to reduce the surface positive charge within the identified charge patches. Substantial reductions in non-specific endocytosis were measured for all mutants. ASA_hIgG1-WT_, wild type Fc human IgG1 anti-streptavidin antibody; CDR, complementarity-determining region; WT, wild type anti-IL-4Rα mAb; LC, light chain; CDR H1, the first complementarity-determining region of the heavy chain; CDR H3, the third complementarity-determining region of the heavy chain. **D** Reduction in non-specific endocytosis via the EEES mutation significantly increased serum exposure relative to the WT anti-IL-4Rα mAb in Tg276 mice following a 1 mg/kg intravenous bolus dose. N = 3 separate animals for EEES, and n = 2 for the WT anti-IL4Rα mAb due to the detection of anti-drug antibodies in one animal at 264 h. WT anti-IL-4Rα mAb data the same as is reported in Figure 1.

**Figure 5:**
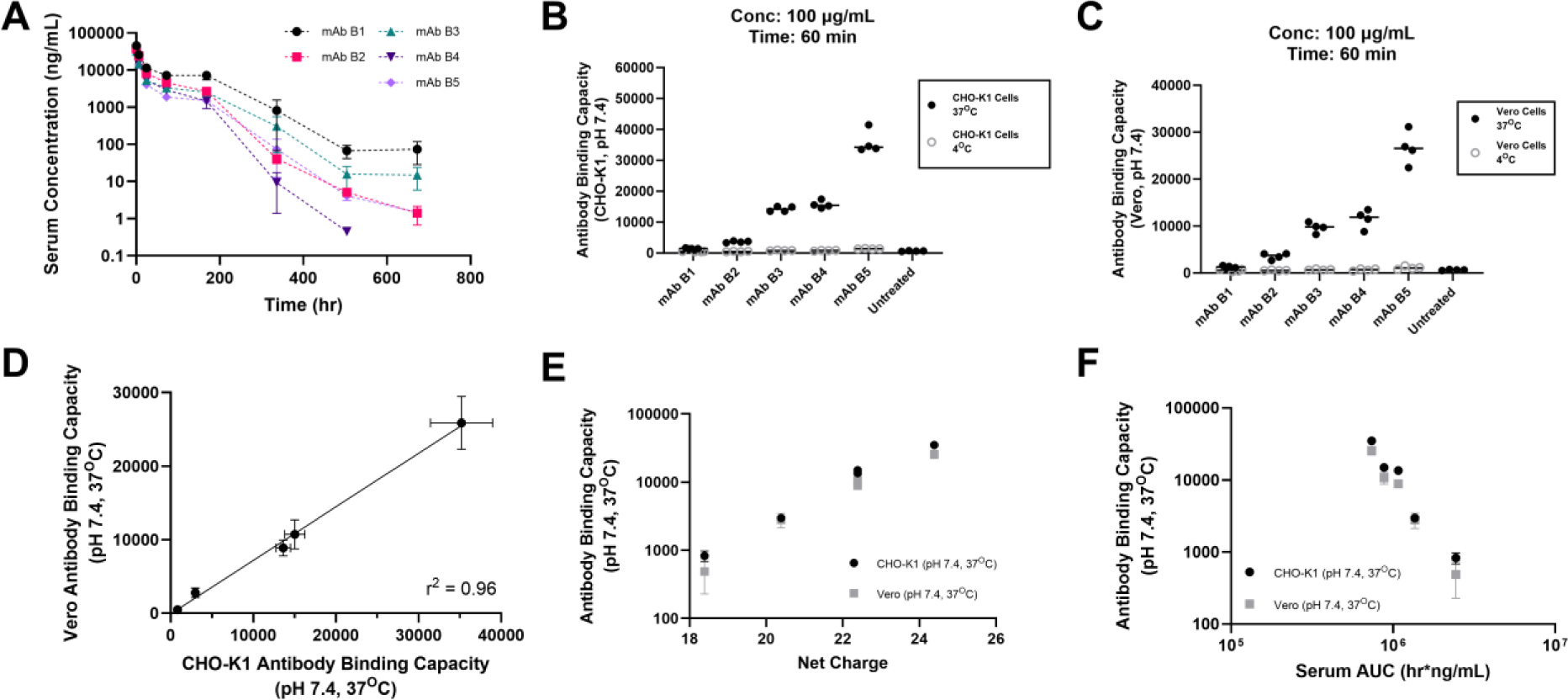
Non-specific endocytosis is conserved across different species and cell types. **A** Serum concentration-time profiles of the preclinical mAbs B1-B5 following single intravenous bolus dose in BALB/c mice. The antibodies exhibited an array of serum exposures. **B, C** Uptake studies were conducted in either (**B**) CHO-K1 (*Cricetulus griseus*, ovarian epithelial-like) or (**C**) Vero cells (*Cercopithecus aethiops*, kidney epithelial) using a mAb panel possessing varying dispositions in wild type BALB/c mice. A range of uptake extents was observed. These results also demonstrated the superior sensitivity of the endocytosis assay performed at 37°C when compared to cell surface binding alone (i.e. 4°C groups). (**D**) Results in Vero and CHO-K1 cells strongly corresponded with each other, supporting non-specific endocytosis as the mechanism for mAb internalization. (**E**) Net charges of the panel B mAbs were found to correlate with NSE rates in Vero and CHO-K1 cells, with more positive net charges leading to higher NSE. (**F**) Additionally, NSE into either CHO-K1 or Vero cells correlated to serum exposures, indicating NSE as an important attribute for the disposition of the tested mAbs. All data plotted as mean ± SD, n = 3-4 per group. CL, clearance; CHO-K1, Chinese Hamster Ovary cells; AUC, area under the curve; NSE, non-specific endocytosis.

**Figure 6:**
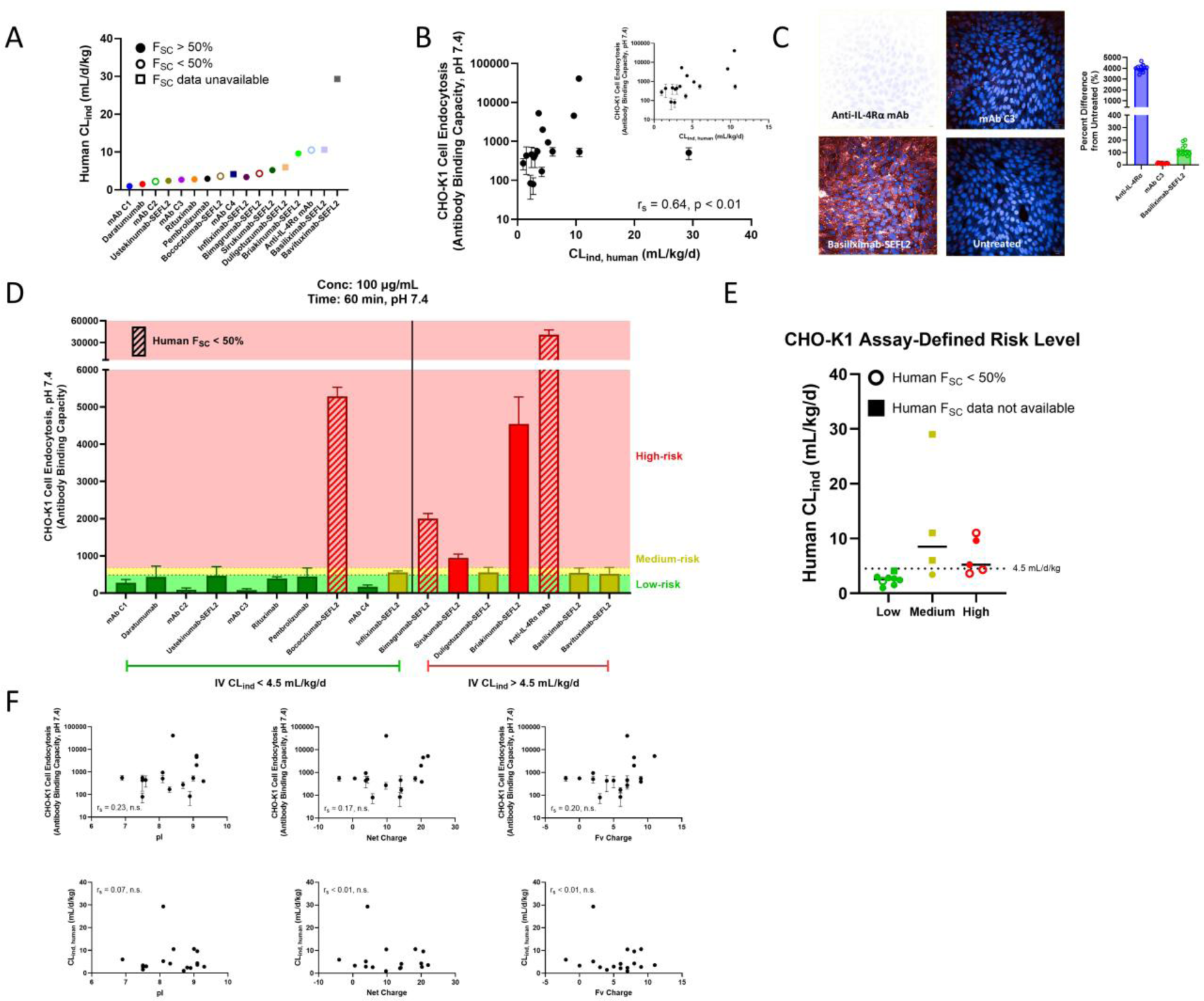
High Non-Specific Endocytosis as an Indication for Suboptimal Clinical Pharmacokinetics. **A** Seventeen mAbs with human PK data were obtained which possessed estimates for CL_ind_. From these, 75% contained F_SC_ estimates in human. **B** Antibodies were analyzed using the CHO-K1 assay as described in Figure 3 to obtain cellular antibody binding capacities (ABCs) and a range of behavior was measured. A relationship was observed between non-specific endocytosis and CL_ind_ in humans with a Spearman correlation (r_s_) of 0.62, p < 0.05. Inset image has removed bavituximab to allow for better visualization of the other mAbs. **C** Confocal imaging on selected mAbs confirm results using the flow cytometry method and exemplify the plasticity of the current approach to use a variety of detection techniques. Scale bar 20 μm, 40X objective. **D** Assay thresholds were generated to better interpret the data. The “Low-risk” group contains mAbs with low-risk of high CL_ind_ and/or low F_SC_ (due to non-specific endocytosis). It was defined as the upper bound of the 95% confidence interval of the mean for all mAbs with F_SC_ above 50% and/or CL_ind_ < 4.5 mL/d/kg. The “Medium-risk” group contains mAbs that may or may not exhibit high CL_ind_ and/or low F_SC_ and was set as two standard deviations above the mean ABC mentioned directly above. The “High-risk” group highlights mAbs with obvious non-specific endocytosis that will likely be detrimental to their disposition in human. The mAbs were then grouped by their performance in the CHO-K1 assay (i.e. low, medium, or high risk) and plotted against their respective CL_ind_ values. **E** demonstrates that an increased risk as defined via higher rates of non-specific endocytosis was an indication for not only higher CL_ind_ following IV administration, but also decreased F_SC_ in human. Of note, every mAb characterized as high-risk exhibited CL_ind_ in human above 4.5 mL/kg/d, F_SC_ below 50%, or both. **F** Relationships were evaluated between pI, net charge, and Fv charge with CHO-K1 endocytosis amounts (upper graphs) or human CL_ind_ (lower graphs). No significant Spearman correlations were observed.

### Homology Modeling and Anti-IL-4Rα mAb Mutant Design

Antibody homology models were built using Molecular Operating Environment (MOE v2022.02, Chemical Computing Group; Montreal, Canada) and default settings in the Antibody modeler (Amber10 forcefield, template search by identity, highest scoring template used in model, number of models = 1). For each antibody, the Fv region was modeled using templates of highest identity for framework and complementarity-determining regions (CDRs). Following structure template searching of the PDB database by framework and CDR regions similarity and identity (Kabat definitions), the highest scored structure template was used for homology modeling. One homology model was then built for each antibody. Structure preparation was performed using default settings for charge, protonation, rotomer and steric clash minimization, and energy minimization. On the prepared homology models, protein properties were calculated at pH 7.4. Protein properties of focus included Fv charge separation (between heavy and light domains), isoelectric point (pI, structure based), zeta potential, dipole moment, hydrophobicity moment, and mobility. Protein patches larger than 50 Angstrom area (hydrophobic, positive, and negative) were also calculated and visualized. Homology models are represented as ribbon structures with surface patches colored by hydrophobic in green, positive in blue, and negative in red.

Comparison of the anti-IL-4Rα mAb and ASA_hIgG1-WT_ homology models and calculated properties highlighted the high positive charge on the anti-IL-4Rα mAb Fv (+7) and resultant higher pI (9). Targeted mutant designs were generated to reduce charge patches and charge separation between variable domains. Specifically, residues in positive charge patches were mutated to negative and neutral residues and protein attributes were calculated to evaluate the effects on charge patch (reduced size) and protein properties (reduced Fv charge and pI). Triple mutants showed the largest effect on Fv charge and pI reduction.

### Compilation of mAbs with Clinical PK Data

PK information was obtained from internal Amgen clinical reports, regulatory body monographs, and/or published literature (14, 15, 18–27). CL_ind_ following a single intravenous bolus dose was defined from these sources as either: (1) clearance obtained via non-compartmental analysis at target saturating doses (i.e. linear PK present; clearance constant with increasing dose) in which we opted to utilize the highest dose data available, (2) estimates of CL_ind_ as determined by Grinsphun et al. (15), or (3) the linear clearance estimate from compartmental modelling with either linear clearance from the central compartment alone or parallel linear and non-linear clearance pathways (e.g. quasi steady-state approximation of the target mediated drug disposition model (14)). Due to the lack of published reports on the clinical disposition of briakinumab, its CL_ind_ was estimated via reported volume of distribution (V_D_) and terminal half-life (t_1/2β_) values obtained under linear PK conditions using the equation below (24, 28, 29):

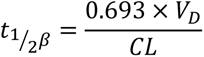

An average body weight of 80 kg was used to normalize the CL_ind_ values. For F_SC,_ a preference was made for absolute subcutaneous bioavailability values in which the parameter was calculated by the ratio of the area under the curve (AUC) for the subcutaneous and intravenous doses. If matching doses were not available, the highest doses were incorporated into the following equation:

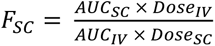

If neither of these were performed, bioavailability estimates from computational modeling were reported, such as from Kakkar et al.(14). Bioavailability ranges were included where specific estimates could not be ascertained from the indicated references.

### Calculation of Physicochemical Properties

Whole-protein net charge, pI, and Fv net charge were calculated utilizing the mAb amino acid sequences at pH 7.4 (30). Charge calculations utilized the Henderson-Hasselbalch equation to compute the contribution of each ionizable region from the pKa in which the fraction of acidic sites deprotonated was 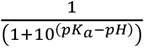 and the fraction of basic sites protonated being 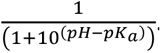. Net or Fc charge was then obtained via

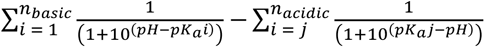

where *n_basic_/n_acidic_* was the number of basic/acidic sites and *pK_a_i/pK_a_j* were the *pK_a_* values from each basic/acidic location, respectively.

## Results

### ASA_hIgG1-WT_ and Anti-IL-4Rα mAb Possess Varying Degrees of CL_ind_ and NSE in Mice

The utilized anti-IL-4Rα mAb is an older Amgen therapeutic protein which lacked mouse PK data due to the use of a murine surrogate mAb for previous preclinical rodent studies. To better understand the mechanisms driving the high rate of its CL_ind_, PK studies were performed in human FcRn transgenic mice (Tg276) to compare the non-target mediated elimination of the anti-IL-4Rα mAb with a reference mAb, ASA_hIgG1-WT_ (31). Single dose PK studies in humanized FcRn Tg276 mice demonstrated the anti-IL-4Rα mAb possessed substantially reduced serum exposure compared to the ASA_hIgG1-WT_ (**Table 1**, **Figure 1**). This result was likely not due to the difference in IgG subclass between the mAbs as past studies have reported no significant effect of IgG subclass on mAb PK (15, 32). Rather, the observations made in Tg276 mice indicated a role for CL_ind_ based on the absence of target, which is due to negligible cross-reactivity with murine IL-4Rα (data not shown).

**Table 1:** Human and rodent CL values for preclinical mAbs obtained via non-compartmental analysis. AUC, area under the curve; CL, clearance; ASA_hIgG1-WT_, wild type Fc human IgG1 anti-streptavidin antibody. An average body weight of 25 g was used for assessments. ASA_hIgG1-WT_, anti-IL-4Rα mAbs, and the mAb B panel data were obtained from three separate *in vivo* studies.

| **mAb** | **Serum AUC**<br>**(hr\*ng/mL)** | **CL**<br>**(mL/hr)** | **Dose**<br>**(mg/kg)** | **Mouse**<br>**Strain** |
| --- | --- | --- | --- | --- |
| ASA<sub>hlgG1-WT</sub> | 2.32E+07<br>(1.57E+07) | 1.39E-03<br>(7.09E-04) | 1 | Tg276 |
| Anti-IL-4Rα mAb WT | 2.77E+05<br>(1.63E+04) | 9.04E-02<br>(5.31E-03) | 1 | Tg276 |
| Anti-IL-4Rα mAb EEES | 6.68E+06<br>(2.58E+05) | 3.75E-03<br>(1.42E-04) | 1 | Tg276 |
| mAb B1 | 2.45E+06<br>(5.42E+05) | 2.10E-02<br>(4.15E-03) | 2 | BALB/c |
| mAB B2 | 1.36E+06<br>(1.32E+05) | 3.69E-02<br>(3.41E-03) | 2 | BALB/c |
| mAb B3 | 1.08E+06<br>(8.70E+04) | 4.65E-02<br>(3.58E-03) | 2 | BALB/c |
| mAB B4 | 8.82E+05<br>(1.76E+05) | 5.82E-02<br>(1.11E-02) | 2 | BALB/c |
| mAb B5 | 7.39E+05<br>(7.11E+04) | 6.81E-02<br>(6.22E-03) | 2 | BALB/c |

A previous analysis of the anti-IL-4Rα mAb in Phase 1 and 2 studies showed non-dose proportional plasma exposure indicative of possible target-mediated clearance dynamics within the range of doses investigated. The clinical results were mathematically described via a two-compartment population PK model with parallel linear (target-independent) and non-linear (target-mediated) clearance pathways (14). From this report, the anti-IL-4Rα mAb was shown to display an estimated CL_ind_ of 11 mL/d/kg, which is more than twice the average CL_ind_ value for the 64 clinical mAbs evaluated by Grinshpun et al. (4.84 ± 4.72 SD) (14, 15). This CL_ind_ was also significantly higher when compared to that of dupilumab, a human IgG4 monoclonal antibody possessing the same antigen (human IL-4Rα) (33). When taken together, these findings indicate target-mediated disposition was not the sole reason for the high CL_ind_ of the previous Amgen anti-IL-4Rα mAb in humans.

To determine if NSE was contributing to the rapid CL_ind_ of the anti-IL-4Rα mAb, internalization kinetics of it and the reference mAb ASA_hIgG1-WT_ were conducted using parental CHO-K1 cells (**Figure 2A**). This line was chosen because it is highly amenable to cell-based assay development, easy to grow, and non-human. Of the two mAbs tested, the anti-IL-4Rα mAb displayed pronounced endocytosis with biphasic time-dependent (**Figure 2B**) and linear concentration-dependent uptake (**Figure 2C**) under the conditions tested. ASA_hIgG1-WT_ was nearly undiscernible from untreated controls such that both populations exhibited strong overlap (exemplified in **Figure 2A**). Additionally, very low signal was observed for the anti-IL-4Rα mAb at 4°C relative to the amount detected at 37°C that indicated the degree of internalization at 37°C greatly outpaced the amount bound at the cell surface.

The ASA_hIgG1-WT_ (hIgG1) and anti-IL-4Rα mAb (hIgG2) were on separate IgG subclasses. To confirm internalization differences were not caused by this, we performed an endocytosis study on anti-IL-4Rα mAb, ASA_hIgG1-WT_, ASA_hIgG2-WT_ to match the anti-IL-4Rα mAb subclass, and ASA_hIgG1-SEFL2_ to evaluate a mutated hIgG1 Fc region lacking effector function (16). No differences were measured for any of the ASA mAb variants (**Figure 2D**). Confocal microscopy performed following a separate 60 min uptake study at 37°C using 100 µg/mL of ASA_hIgG1-WT_ or anti-IL-4Rα mAb confirmed high internalization of the anti-IL-4Rα mAb when compared to ASA_hIgG1-WT_ and untreated controls (**Figure 2E**). These combined observations supported extensive NSE of the anti-IL-4Rα mAb relative to all tested ASA mAbs in CHO-K1 cells (9, 10). Furthermore, NSE of the evaluated ASA mAbs was nearly undetectable when compared to untreated controls. These measurements highlight a quantifiable liability that we propose was a contributing factor for the rapid elimination of the anti-IL-4Rα mAb *in vivo*.

### Development of a Quantitative Flow Cytometry Method to Enable between Day Comparisons

One of the limitations of the initial endocytosis assay iteration was the inability to compare the extent of uptake across experimental days. This was due to the readout being median fluorescent intensity, which is a variable impacted by an assortment of factors that include different voltage settings on the cytometer and using separate instruments across days. To standardize our assays to permit cross-day comparisons and reliable databasing of molecule attributes, we optimized a quantitative flow cytometry method using commercially available anti-mouse IgG microspheres. Each population of these beads binds a pre-determined amount of antibody that can be used to compute the antibody binding capacity (ABC) of a cell population (i.e., the amount of antibody associated with a single live cell event). We began by first ensuring each bead standard was saturated under the experimental conditions. Histograms for each bead population overlapped when stained with either 5 or 10 µg/mL of an anti-human Fc mouse mAb conjugated with Alexa Fluor 647 (**Figure 3A**). Both concentrations resulted in standard curves with similar extents of linearity (r^2^ = 0.999).

We evaluated assay reproducibility by conducting endocytosis studies using anti-IL-4Rα mAb or ASA_hIgG1-WT_ over three separate days, two different cytometers, and two different scientists (n = 20 total samples). Antibodies were incubated with cells at 37°C for 60 min at 100 µg/mL based on our initial kinetic characterization (**Figure 2**). A 60 min incubation time was chosen to maximize sensitivity and broaden our dynamic range of detection to encompass mAbs with more modest NSE rates. Under these conditions, we measured a mean ABC of 40,839 (SD 6614, CV 16.2%) for anti-IL-4Rα mAb and a mean ABC of 921 (SD 315, CV 34.2%) for ASA_hIgG1-WT_ (**Figure 3B**). Population histograms were included for representative ASA_hIgG1-WT_ and untreated samples to demonstrate the significant amount of signal overlap, signifying the weak internalization of ASA_hIgG1-WT_ in CHO-K1 cells (**Figure 3C**). A schematic overview of the assay is included in **Figure 3D**.

### High NSE is Driven by Charge-Dependent Interactions and is Conserved Across Species and Cell Types

One likely mechanism leading to the high rates of anti-IL-4Rα mAb NSE is localized charge in solvent accessible regions of the mAb variable domains. Differences in protein charge have been demonstrated to influence mAb PK, with positive and negative net charge extremes typically associated with the strongest impacts (4, 5, 34, 35). It has been shown that excess localized charge in in the CDRs can affect non-specific mAb endocytosis, even without a large shift in isoelectric point (pI) (5). To provide a better understanding of the relationship between these variables and NSE, molecular descriptors of the charge distribution of anti-IL-4Rα mAb and ASA_hIgG1-WT_ were calculated based on structural modeling (**Figure 4A, B**). The pI of anti-IL-4Rα mAb and ASA_hIgG1-WT_ were calculated from their amino acid sequences as 8.8 and 7.2, respectively. Antibody homology modeling identified several charge patches on the anti-IL-4Rα mAb, including two positive patches in heavy chain CDRs. The anti-IL-4Rα mAb also had a higher calculated charge within the Fv (+7) compared with ASA_hIgG1-WT_ Fv (+2).

The extra positive charge patches on the anti-IL-4Rα mAb together with its higher Fv charge may have contributed to high NSE and CL_ind_. To test this hypothesis, anti-IL-4Rα mAb point mutants were constructed by modifying the CDRs and associated light chain framework regions of the light chains in an attempt to mitigate the parent anti-IL4Rα mAb positive charge attributes. Specific mutations were made at amino acid positions that contributed to clusters of positive charge and/or those critical for driving the net charge of the protein (**Table 2**). ABC values obtained via endocytosis assays in CHO-K1 cells demonstrated each mutation resulted in a substantial reduction in NSE (**Figure 4C**). Single-dose studies in Tg276 mice confirmed that reduction of NSE via the EEES mutations resulted in significantly elevated serum exposure (**Figure 4D**, **Table 1**). ADA assessments identified immunogenicity at 264 h in one mouse that received WT anti-IL-4Rα mAb and was therefore omitted from the plots. These findings indicate positive charge exposed at the surface of the anti-IL-4Rα mAb was a key factor contributing to the elevated rates of NSE and CL_ind_.

**Table 2:** Series of anti-IL-4Rα mAb point mutants to reduce either positive charge patches or the net charge of the protein. Bolded letters correspond to mutant labels within Figures. VH, variable heavy chain; VL, variable light chain.

| Antibody | Mutations | pI | Net Charge |
| --- | --- | --- | --- |
| Anti-IL-4Rα mAb | None | 8.8 | 9.8 |
| Anti-IL-4Rα mAb-SSL | VH:R20S_R33S_R110L | 8.2 | 3.8 |
| Anti-IL-4Rα mAb-SSLS | VH:R20S_R33S_R110L,VL:R95S | 7.8 | 1.8 |
| Anti-IL-4Rα mAb-SELS | VH:R20S_R33E_R110L,VL:R95S | 7.3 | -0.16 |
| Anti-IL-4Rα mAb-SEES | VH:R20S_R33E_R110E,VL:R95S | 7.1 | -2.2 |
| Anti-IL-4Rα mAb-EEES | VH:R20E_R33E_R110E,VL:R95S | 6.8 | -4.2 |

We next performed NSE assessments for a separate panel of preclinical mAbs with a shared target that possessed similar pI values and an expected inability to bind to murine antigen, but dramatically different CL_ind_ in wild type mice (**Table 3**, **Figure 5A**). The mAbs differed at amino acids 27 (light chain), 54, and 56 (both on the heavy chain) within their CDRs. This variance was anticipated to lead to modifications in net and local charge due to the physicochemical characteristics of the amino acids (**Table 3**). CHO-K1 endocytosis results aligned with this trend where mAb B1 exhibited the lowest ABC and mAb B5 the highest (**Figure 5B**). Experiments repeated in green monkey kidney epithelial cells (Vero cells, *Cercopithecus aethiops*), a cell unrelated to CHO-K1 in species (*Cricetulus griseus*) and tissue origin, agreed with results using CHO-K1, which confirmed internalization was non-specific (**Figure 5C, D**). Additionally, uptake studies performed at 4°C supported the lack of target expression in either cell line and also highlighted the magnitude of NSE relative to cell surface binding. We next confirmed the relationship between higher positive net charge and NSE for mAb panel B (**Figure 5E**). *In vivo-in vitro* correlations were then conducted using ABC readouts from either CHO-K1 or Vero cells at 37°C, which demonstrated a strong relationship with serum exposure in wild type mice (**Figure 5F**). Together, these results confirm exposed positive charge at the surface of therapeutic proteins results in higher rates of NSE, reveal NSE patterns are shared across distinctly different cell types, and highlight the utility of direct measures of NSE to inform on CL_ind_.

**Table 3:** Properties of the Preclinical mAb Series. Isoelectric point (pI), net charge, and complementarity-determining region (CDR) amino acid differences of the preclinical mAbs.

| mAb | pI | Net Charge | CDR Sequence Differences<br>Heavy Chain \|\| Light Chain |
| --- | --- | --- | --- |
| mAb B1 | 9.1 | 19 | ISY<u>D</u>G<u>G</u>NKYY \|\| CRAS<u>Q</u>SVGR |
| mAb B2 | 9.1 | 20 | ISY<u>D</u>G<u>G</u>NKYY \|\| CRAS<u>K</u>SVGR |
| mAb B3 | 9.2 | 22 | ISY<u>Q</u>G<u>R</u>NKYY \|\| CRAS<u>Q</u>SVQR |
| mAb B4 | 9.2 | 22 | ISY<u>I</u>G<u>G</u>NKYY \|\| CRAS<u>K</u>SVGR |
| mAb B5 | 9.3 | 24 | ISY<u>Q</u>G<u>R</u>NKYY \|\| CRAS<u>K</u>SVGR |

### High NSE as a Risk Factor for Poor Clinical PK Behavior

We sought to expand our findings by directly measuring NSE of biologics with clinical PK data. A total of 17 mAbs were obtained that had varying extents of CL_ind_ estimates in humans (**Figure 6A**, **Table 4**). NSE studies were conducted in CHO-K1 cells as described in **Figure 3** using flow cytometry with the ABC as the assay readout. A relationship was observed between extent of NSE at pH 7.4 over 60 min and CL_ind_ in humans (**Figure 6B**, Spearman correlation coefficient 0.62, p < 0.01). Immunofluorescence with high content confocal microscopy was subsequently performed on select mAbs with increasing NSE to support these observations. Additionally, we aimed to highlight how different experimental and analysis approaches (i.e. not using flow cytometry with ABC interpolations) could still result in the same outcome. We measured by confocal microscopy elevated NSE in CHO-K1 cells for the WT anti-IL-4Rα mAb, basiliximab research analog, and mAb C3 (**Figure 6C**), consistent with results from the flow cytometry assay. Of note, microscope settings differed from those in **Figure 2** to simultaneously capture the internalization dynamics of the NSE behavior for several mAbs within the same experiment. To ensure the detectors were not saturated on the confocal microscopes, fluorescent microsphere curves were generated daily as described in **Figure 3** and imaged under the same experimental settings as the cells. Linearity was observed each day with linear regression fits equaling or exceeding r^2^ values of 0.99 (**Supporting Figure 1**).

**Table 4:**
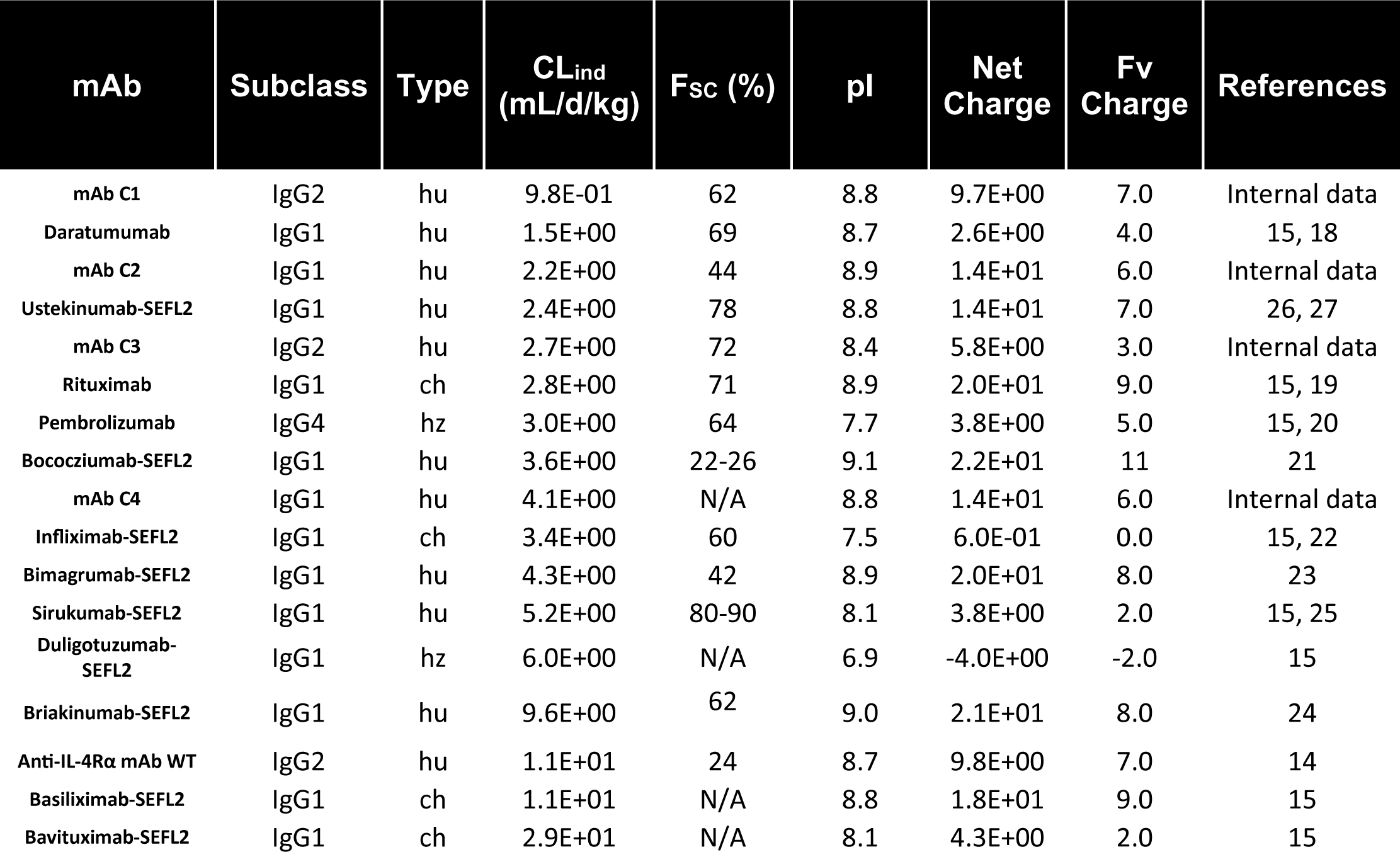
Pharmacokinetic and physicochemical properties of 17 monoclonal antibodies with clinical data. N/A; data unavailable; hu, human; ch, chimeric; hz, humanized.

| **mAb** | **Subclass** | **Type** | **CL<sub>ind</sub>**<br>**(mL/d/kg)** | **F<sub>sc</sub> (%)** | **pI** | **Net**<br>**Charge** | **Fv**<br>**Charge** | **References** |
| --- | --- | --- | --- | --- | --- | --- | --- | --- |
| mAb C1 | IgG2 | hu | 9.8E-01 | 62 | 8.8 | 9.7E+00 | 7.0 | Internal data |
| Daratumumab | IgG1 | hu | 1.5E+00 | 69 | 8.7 | 2.6E+00 | 4.0 | 15, 18 |
| mAb C2 | IgG1 | hu | 2.2E+00 | 44 | 8.9 | 1.4E+01 | 6.0 | Internal data |
| Ustekinumab-SEFL2 | IgG1 | hu | 2.4E+00 | 78 | 8.8 | 1.4E+01 | 7.0 | 26, 27 |
| mAb C3 | IgG2 | hu | 2.7E+00 | 72 | 8.4 | 5.8E+00 | 3.0 | Internal data |
| Rituximab | IgG1 | ch | 2.8E+00 | 71 | 8.9 | 2.0E+01 | 9.0 | 15, 19 |
| Pembrolizumab | IgG4 | hz | 3.0E+00 | 64 | 7.7 | 3.8E+00 | 5.0 | 15, 20 |
| Bococizumab-SEFL2 | IgG1 | hu | 3.6E+00 | 22-26 | 9.1 | 2.2E+01 | 11 | 21 |
| mAb C4 | IgG1 | hu | 4.1E+00 | N/A | 8.8 | 1.4E+01 | 6.0 | Internal data |
| Infliximab-SEFL2 | IgG1 | ch | 3.4E+00 | 60 | 7.5 | 6.0E-01 | 0.0 | 15, 22 |
| Bimagrumab-SEFL2 | IgG1 | hu | 4.3E+00 | 42 | 8.9 | 2.0E+01 | 8.0 | 23 |
| Sirukumab-SEFL2 | IgG1 | hu | 5.2E+00 | 80-90 | 8.1 | 3.8E+00 | 2.0 | 15, 25 |
| Duligotuzumab-SEFL2 | IgG1 | hz | 6.0E+00 | N/A | 6.9 | -4.0E+00 | -2.0 | 15 |
| Briakinumab-SEFL2 | IgG1 | hu | 9.6E+00 | 62 | 9.0 | 2.1E+01 | 8.0 | 24 |
| Anti-IL-4Rα mAb WT | IgG2 | hu | 1.1E+01 | 24 | 8.7 | 9.8E+00 | 7.0 | 14 |
| Basiliximab-SEFL2 | IgG1 | ch | 1.1E+01 | N/A | 8.8 | 1.8E+01 | 9.0 | 15 |
| Bavituximab-SEFL2 | IgG1 | ch | 2.9E+01 | N/A | 8.1 | 4.3E+00 | 2.0 | 15 |

In addition to CL_ind_ estimates, the majority of the mAbs in the clinical panel possessed subcutaneous bioavailability (F_sc_) data in humans (**Figure 6A**). Subcutaneous administration remains the preferred injection route for therapeutic proteins due to added patient convenience. However, subcutaneously administered drugs exhibit variable absorption, often leading to sub-optimal bioavailability, F_sc_. F_sc_ is dependent upon several factors that includes temperature, solubility, lymph flow at the injection site, and tissue composition (36). Past work has related elevated non-specificity to low F_SC_, but to our knowledge, no study has directly connected cellular NSE with F_SC_ (37, 38). It was hypothesized that the factors driving high NSE and elevated CL_ind_ following an intravenous dose would also be important determinants for F_SC_ (39, 40). This is because high rates of NSE at the injection site and/or within the lymphatics could result in elevated therapeutic protein catabolism, leading to less drug available for absorption into the circulation. Additionally, the mechanisms driving high NSE may also lead to other *in vivo* characteristics that are detrimental to F_SC_, such as non-specific interactions with the subcutaneous extracellular matrix (35). We hypothesized that a cell-based measure of NSE might be correlated to mAb F_sc_.

We established three categories of NSE using the flow cytometry data and descriptors of the well-behaved mAbs to provide a generalized example for how cellular NSE data could be used to better understand the disposition of the mAbs. The “Low-risk” group contained mAbs with a low likelihood of high CL_ind_ and/or low F_SC_ (due to NSE). It was defined as the upper bound of the 95% confidence interval of the mean for all mAbs with F_SC_ above 50% (where data was present) and CL_ind_ below 4.5 mL/d/kg, which equates to a half-life of approximately 10 days in an 80 kg individual assuming a volume of distribution (V_D_) of 5 L 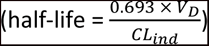(29). The “Medium-risk” group contained mAbs that may or may not exhibit high CL_ind_ and/or low F_SC_ and was set with its upper bound as two standard deviations above the mean ABC mentioned above. The “High-risk” group was anything above the “Medium-risk” cutoff and highlighted mAbs with obvious NSE that will likely be detrimental to their disposition in human. **Figure 6D** illustrates the distribution of mAbs within these bins, aligned left to right with increasing CL_ind_ estimates. The mAbs were then grouped by their performance in the CHO-K1 assay (i.e. low-, medium-, or high-risk) and plotted against their respective CL_ind_ values. **Figure 6E** demonstrates that an increased risk as defined via higher rates of NSE was an indication for not only higher CL_ind_ following IV administration, but also decreased F_SC_ in human. Of note, every mAb characterized as high-risk exhibited CL_ind_ in human above 4.5 mL/kg/d, F_SC_ below 50%, or both.

The preclinical mAbs analyzed within the current work demonstrated trends of reduced total net charge and/or pI with cellular NSE. However, these comparisons were restricted to point mutants of parental mAbs. We examined the relationships between pI, net charge, and Fv charge with NSE within the clinical mAb series to take advantage of more broad physicochemical behaviors. No significant trends were observed between any of the three parameters and CHO-K1 ABC values (**Figure 6F, top**). Furthermore, no association was found between pI, net charge, or Fv charge with CL_ind_ in humans (**Figure 6F, bottom**). These combined results demonstrate the applicability of direct NSE measurements in cells to inform on *in vivo* disposition of mAbs.

## Discussion

Several *in vitro* techniques have been reported to assess the potential of mAb non-specific properties and NSE (3, 13). However, direct measurements of mAb NSE using cells is not routinely conducted for the purposes of evaluating determinants of mAb PK. We have shown in the current report that separate mAbs can possess distinct rates of NSE into mammalian cells due to positive charge on the protein surface, supporting several studies from other research groups (4, 5). Additionally, we generated a reproducible cell-based assay capable of directly quantifying mAb NSE whose general workflow can be readily integrated into *in vitro* ADME screening to triage biologics on the basis of suspected aberrant PK behaviors.

Our detailed flow cytometry method highlighted a specific approach that could be incorporated by others to quantify intracellular protein accumulation. However, a primary aim of our work was not the development of an explicit method but rather to demonstrate the feasibility and utility of direct measurements of NSE to inform on the CL_ind_ of biologics. Flow cytometry was just one of many possible quantitation approaches that could have been pursued, with alternatives including enzyme-linked immunosorbent assay (ELISA), liquid chromatography with tandem mass spectrometry, or confocal microscopy. Each of these assays have their own advantages and limitations. ELISAs would enable full quantitation of intracellular protein amounts (e.g. amount of analyte per amount of total protein in the lysate) but would require efficient and thorough cell lysis and individual standard curves for each test article. High content confocal microscopy, as was utilized here, offers a technique to analyze hundreds to potentially thousands of proteins per day with the incorporation of automation. But it is limited to arbitrary fluorescence unit readouts. The adaptability of the analysis method permits both the incorporation of our overall approach at various stages of drug development as well as the use of the assay for a variety of goals that could range from candidate screening to quantitative measurements of NSE.

The characterization of ASA_hIgG1-WT_ and the anti-IL-4Rα mAb support NSE of a mAb in a cell-based experimental platform lacking target for several reasons. The first was the linear relationship between the concentration of the anti-IL-4Rα mAb and the extent of its endocytosis. Second, we observed a large discrepancy between anti-IL-4Rα mAb cell-associated amounts at the separate temperatures. Whereas 37°C allowed for cell binding, endocytosis, and trafficking, 4°C permitted only surface binding. The low signal at 4°C thus indicated anti-IL-4Rα mAb internalization at 37°C as well as supported our conclusion for the absence of target within CHO-K1 cells. Of note was the biphasic time-dependent uptake of the anti-IL-4Rα mAb. One potential cause of the inflection could be that lysosomal degradation becomes significant during this timeframe, thereby altering the net accumulation rate of detectable antibody. This would be due to detection in the current assay via the anti-human Fc antibody rather than a direct conjugation approach such as labeling with pH-insensitive fluorophores, whose intracellular accumulation would be unaffected by catabolism. A second possibility may have been rapid and constitutive analyte exocytosis following initial internalization, as measured by others (10, 41–43).

The current technique provides an approach that takes advantage of a living cell and intact cell membrane instead of isolated components or surrogates. To strengthen conclusions, modifications can be made such as co-culturing two distinct and unrelated cell types. If flow cytometry and/or confocal microscopy is used for quantification, stably expressed fluorescent reporters (e.g. green fluorescent protein) could easily differentiate between two or more cell types within the same sample. Additionally, readouts from the current assay can directly yield analyte-specific, single-cell internalization rates that can be incorporated into computational models. For example, cellular measures can provide PK modeling efforts with NSE rates and/or rate constants for each biologic of interest rather than inferring this parameter with biophysical techniques or from general measures of fluid-phase endocytosis (44, 45). Furthermore, specific cell types can be used to tailor the internalization kinetic measurements to the tissue being modeled, such as primary human endothelial cells or interstitial resident immune cells. This can provide research groups with a more representative level of site-specific information to assist in a deeper, more biologically relevant assessment of preclinical compounds.

Despite these advantages, there were also limitations in the current study. The preclinical mAb B panel exhibited non-linearity in the terminal phases. Neither target engagement to murine antigen based on the tissue expression and binding behavior of the parental mAb (data not shown) nor anti-drug antibodies were believed to be responsible. For this reason, serum exposure was used to correlate the data, which we showed trended strongly with NSE. Additionally, the set of mAbs with clinical PK data provided an initial dataset to exhibit the capabilities of the currently disclosed method. More proteins and non-mAb modalities (e.g. bispecific antibodies) with diverse *in vivo* behaviors will be needed to expand and confirm the current results and strengthen the relationships identified for cellular NSE with CL_ind_ and F_SC_. This would also offer greater insight into how various physicochemical parameters impact NSE beyond the assessments of pI, net charge, and Fv charge initiated here, which could include an exploration of possible interplays between hydrophobicity, aromaticity, and charge-based features with NSE rates.

Aside from NSE, FcRn-facilitated recycling is another major component of mAb CL_ind_ (1, 2) that was not addressed within the current work. FcRn is crucial for the long half-lives of Fc-bearing therapeutics due to its pH-dependent association with the Fc region. Following endocytosis, FcRn binds to its ligands during endosomal acidification and then mediates ligand trafficking to the plasma membrane where dissociation from FcRn occurs due to the low affinity of the FcRn-Fc interaction at near neutral pH (7). Though FcRn is a vital aspect of mAb PK and variations in the preclinical mAb CL_ind_ could possibly be caused by differences in murine FcRn binding, our efforts were focused on determining the relationship between NSE and CL_ind_. Previous work has demonstrated that a combinatorial evaluation of non-specific and FcRn interactions for mAbs can provide additional insight into potential PK shortcomings (3). Ongoing work within our laboratory is examining these relationships utilizing mammalian cells but were beyond the scope of the current work (subsequent manuscript submitted).

## Conclusion

We have generated a novel cell-based method to quantify NSE of protein therapeutics and successfully employed the assay to determine an anti-IL-4Rα mAb possessed substantial NSE which contributed to its high preclinical CL_ind_. Additionally, we observed a correlation between NSE in CHO-K1 cells and clinical CL_ind_ for a series of mAbs. Because many variables can influence mAb PK, a cell-based measure of NSE should be used in conjunction with accompanying approaches to assist in preclinical candidate selection and de-risking strategies as reported previously (46). Future work to develop amino acid sequence and structure-based predictive models informed by cellular outputs such as these could be of high value to limit late-stage therapeutic attrition. These types of combinatorial strategies will be of the upmost importance as the pharmaceutical industry broadens the structural basis of its development portfolio and looks to more complex therapeutic modalities for treating disease, such as multi-specific antibodies (47).

## Acknowledgements

We thank Jacky Li, Patrick Cannon, Marisela Killian, and Christopher Vandevert from the Amgen San Francisco flow core for their extensive technical support, Marissa Mock for her contribution to the design and facilitation of the preclinical mAb series, and Benjamin Alba for the generation of the ASA_hIgG1-SEFL2_. Additionally, we thank Dan Rock, Joshua Pearson, and Dharam Thakkar for their previous contributions with the mouse PK studies.

## Author Contributions

Experimental design, data acquisition, and/or interpretation of results was performed by all authors; drafting of the final manuscript (M.A.B., M.T.H.T., C.D.S., K.D.C., K.P.C.); all authors have approved the final version of this manuscript and agreed to the accuracy of its contents.

---

**Supporting Figure 1:**
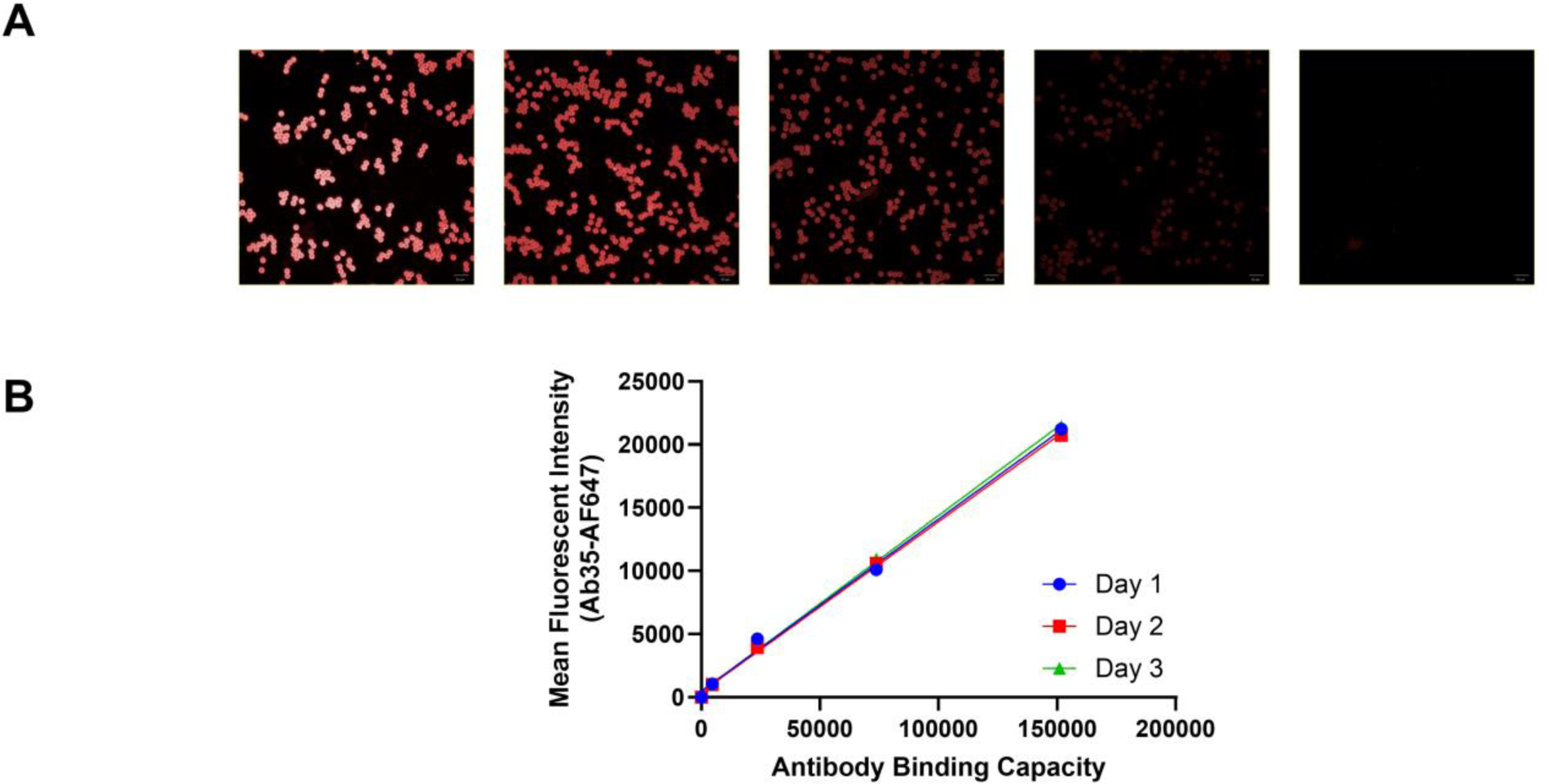
Quantum simply cellular beads were stained daily as described for Figure 3 and imaged alongside CHO-K1 uptake samples. **A** Representative image from one of the days depicted. **B** Quantification of mean fluorescent intensities from the individual bead populations indicated the image acquisition parameters remained within the linear range of the confocal microscope.

